# Modeling brain reorganization after hemispherectomy

**DOI:** 10.1101/2020.12.25.424412

**Authors:** Luis F Seoane, Ricard Solé

## Abstract

Brain reorganization after hemispherectomy (i.e. after the removal of a whole hemisphere) is perhaps the most remarkable example of large-scale brain plasticity. Most often patients survive and recover their skills. Functional traits located in the lost side (e.g. language areas) can sometimes be completely reassembled in the remaining hemisphere, which seamlessly takes on the additional processing burden. This demands drastic rearrangements, perhaps involving the readaptation of functionally and structurally diverse neural structures. We lack mathematical models of how this happens. We introduce a very simple model, based on self-organized maps, that provides a rationale to the clinical aftermath of the intervention, putative windows for recovery, and the origins and nature of observed thresholds for irreversible function loss. The implications for brain symmetry and potential scenarios in simulated pathologies, including efficient suggested treatments, are outlined.

## Introduction

The well-known potential of brains to cope with damage and aging is one of their most defining attributes. Both degradation and regeneration happen on multiple scales, from neurons to brain tissues, and over time. A remarkable example of this regenerative potential is the capacity to withstand brain hemispherectomy (BHEM). This is an extreme (but largely successful) neurosurgical procedure used to treat epileptic disorders that involve severe, life-threatening episodes. BHEM entails removal or disconnection of a whole cerebral hemisphere. Surprisingly, in the ensuing “half a brain” scenario (***Borgstein and Grootendorst, 2002; Devlin et al., 2003; Battro, 2006***) patients can recover cognitive, motor, and somatosensory functions very close to normal (***Pulsifer et al., 2004; Liégeois et al., 2010; Danelli et al., 2013***)—a more striking outcome since subtler unbalances of the connectome result in deeper cognitive impairments (***van den Heuvel and Sporns, 2019***).

The case of language illustrates some challenges after BHEM. In complete brains, language is spontaneously lateralized (most often to the left) following the assembly and maturation of several complex, tightly interlinked cortical regions (***Geschwind, 1972; Galaburda, 1995; Catani et al., 2005; Fedorenko and Kanwisher, 2009; Fedorenko et al., 2012; Fedorenko and Thompson-Schill, 2014; Berwick and Chomsky, 2016***). Children below three who have their left hemisphere removed can still recover language by having these structures reorganize at their right side. Interventions at early age correlate with more successful recovery, while processes later in life most often result in complete language loss. This suggests the existence of *window periods* (around six years of age (***Woods and Teuber, 1978; Woods and Carey, 1979; Marcotte and Morere, 1990; Hertz-Pannier et al., 2002***)) after which function cannot be regained. But evidence is inconclusive, with counterexamples of language recovery at older age showing that the limits are not rigid (***Smith, 1966; Vargha-Khadem et al., 1997; Hertz-Pannier et al., 2002; Voets et al., 2006***).

The successful clinical cases reveal that we can survive and thrive with only one hemisphere, but little is known about how the brain reorganization itself takes place. Two recent fMRI studies registered neural activity from BHEM patients (***Kliemann et al., 2019, 2021***) finding that brain parcellations derived from intact, healthy brains are consistent with activity in hemispherectomized brains. Overall, connections within task-specific (e.g. resting state, motor, attention, theory of mind) networks increase with respect to normal brains. Commonly, connections across modal networks remain as in complete brains. Incidental evidence suggests that more connections between different modal networks (i.e. “misplaced” links) result in worst cognitive performance (***Kliemann et al., 2019***). Thus functional centers appear to take up preferably affine tasks.

Brain reorganization has been proved, among others, by experiments in macaques (***Paul et al., 1972; Merzenich and Kaas, 1982; Merzenich et al., 1983, 1984; MErZENICH et al., 86; Pons et al., 1991***), clinical procedures (***Rivera-Rivera et al., 2017***), and recovery after stroke (***Pascual-Leone et al., 2005***). Simple mathematical models (***Armentrout et al., 1994; Sutton III et al., 1994; Weinrich et al., 1995; Spitzer et al., 1995; Lytton et al., 1999***) illustrate smooth, local processes when topographic maps need to expand (invading adjacent areas) or change their nature slightly. Less consensus emerges at larger scales. Some experiments indicate a spatial limit to plasticity, suggesting that some representations cannot be smoothly transferred to distant parts of a same cortical area (***Merzenich and Kaas, 1982; Merzenich et al., 1983; MErZENICH et al., 86***). Others demonstrate deep rearrangements of topographic maps prompted by drastic transformations of the external anatomy (***Paul et al., 1972; Merzenich et al., 1984; Pons et al., 1991***).

How does this remapping take place? Does it proceed smoothly, or does it need a drastic rewiring of the intact cortical structures? If so, how drastic, and how does it happen? Are new cognitive centers developed, or do existing areas double up their efforts? Is the computational load distributed according to each neural circuit’s specific capabilities, or are the existing parts reconfigured to take on alien functions? Can we capture the complex features and scales involved in BHEM in a simple model?

In this paper we seek to use the minimal key ingredients for brain reorganization to provide a rationale for the clinical and functional features observed after BHEM. We represent higher brain functions abstractly, as the ability of a Self-Organized Map (SOM) (***Kohonen, 1982***) to build globally-coherent and detailed representations out of local input cues. Below, we detail how we model these input stimuli, which might display two conflicting underlying geometries. We simulate BHEM and SOM reorganization under the simplest representations of lateralized stimuli and investigate how this changes as bilaterality becomes increasingly relevant. We further study a priming process (of biological relevance) that enables our SOMs to achieve finer representations. We find that this priming conflicts with strongly bilateral stimuli, resulting in pathological representations. The reorganization of primed SOMs that we observe after simulated hemispherectomy strongly suggests that this intervention can be preferred to fix pathological SOMs—pointing towards unexploited clinical uses of BHEM.

These insights concern mostly somatosensory maps. Next we turn our attention to the language ability, which involves lateralized structures (***Geschwind, 1972; Galaburda, 1995; Catani et al., 2005; Fedorenko and Kanwisher, 2009; Fedorenko et al., 2012; Fedorenko and Thompson-Schill, 2014; Berwick and Chomsky, 2016***) that develop from bilaterally symmetric substrates in newborns (***Galaburda, 1995; Berwick and Chomsky, 2016; Perani et al., 2011***). Therefore we model redundant representations seated at opposing hemispheres, and how they fare before and after BHEM. Our model strongly suggests that lateralized stimuli challenge the inborn bilateral symmetry, forcing the development of a dominant side. Keeping bilaterality risks a deterioration of both candidate representations. We observe how the dominated hemisphere degrades through the lateralization process, and how this degradation is key to explaining window periods within which language recovery is possible—and beyond which it becomes rare. Our results are in accordance with empirical observations and propose a mechanism that explains phenomenology around these window periods.

## Results

We run a series of numerical experiments of simulated hemispherectomy. In each of them, we first train a SOM on either linear or bilateral stimuli (Fig. 5(a)), then remove half of its units (Fig. 5(b)), then retrain the remaining network on stimuli of the same kind. For training, we fix model parameters for the sensory input (*L, σ*_*s*_, and *p*_*T*_) and the SOM (*N, σ*_*t*_, and *p*_*P*_). Then we produce *T*_*t*_ stimuli examples to be presented to the SOM, each peaked at a random, uniformly distributed *x*^0^ ∈ [0, *L*]. We measure the distance of the resulting representation, *u*^0^(*x*^0^), to either the optimal linear, 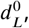, or bilateral, 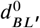, representations. The superindex 0 indicates that this evaluation happens before the

SOM is halved.

To simulate a BHEM, we copy half (either the first or last) contiguous units into a smaller SOM (Fig. 5(b)). We evaluate this halved SOM’s representation right after hemispherectomy, *u*^*h*^(*x*^0^), by measuring the distances 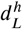 or 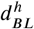. Note that these networks will be missing proper representations for some of the input stimuli. We term these stimuli ‘orphan’. Finally, we retrain the halved SOM for *T*_*rt*_ time steps with new input strings. After re-training, we evaluate the SOM’s representation, *u*^*rt*^(*x*^0^), again with respect to linear, 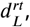, or bilateral, 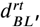, optimal representations.

The ability to establish finely-tuned and globally-coherent representations depends on *σ*_*s*_ and *σ*_*t*_ (Sup. Figs. 2 and 5). If *σ*_*s*_ ≪ *L* and *σ*_*t*_ ≪ *N* (Sup. Fig. 2(a-c)), linear representations form locally, but not globally. As the training processes begins, locally coherent representations start forming, then grow like a crystal as nearby-peaked stimuli are shown as training input. This happens at several locations over the range of SOM units. At each seed there is a choice between the two optimal representations, 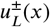. Local choices are random and might collide with others as they expand, resulting in a globally fragmented *u*(*x*^0^). From a somatosensory perspective, this would be equivalent to representing two adjacent patches of skin in distant cortical regions.

Larger *σ*_*s*_ and *σ*_*t*_ result in more global coherence (Sup. Fig. 2(d-f)). But the emerging representations are fairly coarse-grained (compare the distributions of weights in Sup. Fig. 2(b) vs. 2(e)). This means that individual units of these globally-coherent maps are activated by a broad range of stimuli. This corresponds to a somatosensory representation that responds to stimulation of distant, unrelated patches of skin. Introducing *primed* SOMs circumvents this problem.

### Simulated hemispherectomy of unprimed SOMs

We want to study BHEM in relatively good maps, which model somatosensory representations in intact brains. We do not care about fragmented or otherwise pathological maps in his work. It is challenging to obtain optimal linear representations with unprimed SOMs. We explored the model visually to select fair, unprimed SOMs (Sup. Fig. 5).

Sup. Fig. 7 illustrates BHEM in an unprimed (*p*_*P*_ = 0) SOM trained with fully linear stimuli (*p*_*T*_ = 0). Immediately after hemispherectomy, the SOM representation (*u*^*h*^(*x*^0^), light gray in Sup. Fig. 7(a); weights in Sup. Fig. 7(c)) is missing its natural response to half of the stimuli. This “orphan domain” extends from *x*^0^ = 0 to *x*^0^ = *L/*2. SOMs always categorize any input, even if it is new or unexpected. Here, orphan inputs close to the center (*x*^0^ ≲ *L/*2) are parsimoniously mapped to the representation of the first non-orphan stimulus—because Hebbian learning guarantees that a neighborhood of units responds to a neighborhood of inputs. As we move towards the extreme of the orphan domain (*x*^0^ → 0), stimuli become represented at the opposite end of the halved SOM. This is due to the anti-Hebbian mechanism of SOMs (see Methods), which represses the response to moderately distant signals more strongly than to very distant ones. Orphan stimuli at the extreme (*x*^0^ → 0) are actually being assigned not to the most affine units, but to the units that are least indifferent to those orphan signals.

New representations emerge after 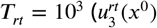, medium gray in Sup. Fig. 7(a)) and 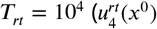, dark gray in Sup. Fig. 7(a)) retraining steps (weights in Sup. Fig. 7(d) and (e) respectively). They grow from the original map, *u*^*h*^(*x*^0^). Intact representations are gradually squeezed into smaller SOM regions to accommodate orphan stimuli (Fig. 3(a)). The process seems smooth, as units at the edge of the (shrinking) orphaned domain are parsimoniously incorporated. This is consistent with earlier studies of lesions in topographic maps based on SOM (***Armentrout et al., 1994; Sutton III et al., 1994; Weinrich et al., 1995; Lytton et al., 1999***).

This parsimonious rearrangement might seem convenient, but it requires the relocation of all intact representations. In neuro-anatomical terms this seems possible to correct small injuries, for which orphaned function can be *stuffed* into relatively similar, adjacent tissue. But after a hemispherectomy this would imply that well-established representations would be displaced far away from their place. Motor control regions must keep track of these changes too. Thus, over large scales, the parsimonious rearrangement appears costly and unlikely.

Things get more interesting with bilateral stimuli. For a fixed set of parameters (see Fig. 2(a-b)) we explored a range of input bilaterality, *p*_*T*_ ∈ [0, 0.5]. Fig. 2(a) shows 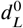 (black) and 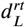 (red). For small *p*_*T*_ (≲ 0.1), the emergent maps ignore the bilateral component and result in mostly linear representations 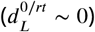. Most halved SOMs, again, reorganize themselves by squeezing orphan stimuli as before (Figs. 2(e) and 3(b)). In average, complete SOMs achieve better maps than those after BHEM 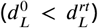. For *p*_*T*_ > 0.1, these linear representations degrade quickly. In turn, maps closer to the optimal bilateral emerge (Fig. 2(b))—however, the resulting map is far from perfect, illustrating the conflict between both geometric patterns underlying input signals. Outstandingly, the halved SOM after retraining shows better bilateral maps than the full original network (Fig. 3(d)). This suggests that BHEM might amend problems emerging on complete SOMs when neither optimal representation can emerge.

### Simulated hemispherectomy of primed SOMs

Unprimed representations are coarse, which results in relatively large 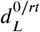 and 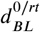. Primed SOMs render globally coherent yet fine-grained maps closer to a lateralized optimum. This mechanism is developmentally justified: phantom limbs in newborns with missing extremities (***Spitzer et al., 1995***) prove that somatosensory maps grow from innate blueprints acquired over evolutionary time.

We explore BHEM in primed SOMs (*p*_*P*_ = 0.5) with fixed parameters (see Fig. 2(c-d)) for the same varying levels of input bilaterality (*p*_*T*_ ∈ [0, 0.5]). Representations in complete SOMs for low *p*_*T*_ (≲ 0.1) turn out much closer to the optimal linear map (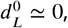, black plot in Fig. 2(c)) than unprimed maps. But, crucially, priming precludes a reorganization into a similarly linear representation after hemispherectomy (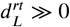for any *p*_*T*_, red plot in Fig. 2(c); Fig. 3(e))—even for stimuli with negligible bilaterality (*p*_*T*_ → 0). Priming has induced a different, sturdy topological attractor in the halved SOM which, now, cannot be reorganized by parsimoniously squeezing orphaned stimuli into the intact substrate. Instead, orphan stimuli are readily mapped into their bilateral counterparts (Figs. 2(f) and 3(f)).

This primed attractor also prevents the complete SOM from ever developing a bilateral representation (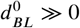 for all *p*_*T*_, black plot in Fig. 2(d)). The interference with the primed map becomes hazardous as the bilateral component increases—for *p*_*T*_ ≳ 0.1 the linear representation is destabilized as well. Thus, an imbalance between the rigidity of inborn representations and excess input bilaterality results in pathological SOMs before hemispherectomy. In turn, again, a striking side effect is that halved SOMs swiftly fold into robust bilateral maps (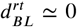 for all *p*_*T*_ except *p*_*T*_ → 0, red plot in Fig. 2(d); Fig. 3(g)).

What does this mean in neuro-anatomical terms? Regarding the impossibility of reorganization into a linear map (Fig. 3(e)): The priming process enables more refined representations (with smaller *σ*_*s*_, *σ*_*t*_), hence signals with close values of *x*^0^ are more dissimilar than before. This hampers the gradual incorporation of orphan stimuli through the map’s edge (opposed to what happened in Sup. Fig. 7). This mechanism works locally at most, in agreement with earlier simulations of local injuries (***Armentrout et al., 1994; Sutton III et al., 1994; Weinrich et al., 1995; Lytton et al., 1999***). On the upper side, stimuli are similar enough to their mirror-symmetric counterparts. This enables the robust folding of halved SOMs into bilateral maps (Fig. 2(f) and 3(f)), which demands less overall reorganization since the intact representations are not dragged across the brain. This offers an insight on why BHEM might work with relatively small cognitive impact. Furthermore, the bilateral maps of halved SOMs appear sturdy even when the linear representations of complete networks break apart (Fig. 3(g)). This puts BHEM forward as a treatment for pathologies in which full, lateralized maps fail to develop. In our model this happens due to an interference between the two optimal topologies: a primed linear geometry versus a strong bilateral component.

### Symmetry breaking of redundant structures—insights for language

The language capacity is housed in several cortical regions around the sylvan fissure of the dominant (usually left) hemisphere (***Geschwind, 1972; Catani et al., 2005; Fedorenko and Kanwisher, 2009; Fedorenko et al., 2012; Fedorenko and Thompson-Schill, 2014; Berwick and Chomsky, 2016***). The same areas in both brain halves start as symmetric at birth (***Galaburda, 1995; Perani et al., 2011; Berwick and Chomsky, 2016***), but only the dominant side matures into a complete language organ. We model two roughly equivalent *left* and *right* SOMs that compete to elaborate a globally coherent representation of fully linear input signals (model parameters in Fig. 4). Each SOM is independently primed with 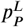 and 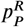 respectively. This ensures that both sides are hinted towards the target representation (i.e. both contain similar precursory circuits), but a slight inter-hemispheric asymmetry is explored by keeping 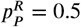 and varying 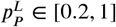.

We train the composed SOM (both sides concatenated), simulate a hemispherectomy, and retrain each independent half. During the initial training, each input has two candidate representations. Symmetry is broken at random but aided by the distinct primings 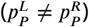. The responding side becomes more apt at representing the shown stimulus, thus amplifying the asymmetry. Fig. 4(a) shows the distance to the optimal representation for left (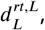, black) and right (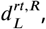, red) SOMs after hemispherectomy and retraining. For very symmetric priming 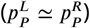 both sides might fail to establish a functioning map, even after individual retraining (Fig. 4(b)). Hence symmetry breaking and the emergence of a dominant side are preferred.

We next executed BHEM after a varying training time *T*_*t*_, modeling surgery at different ages, making ‘left’ the dominant side. The left SOM achieves consistently a good representation (black plot in Fig. 4(c)). The right SOM’s map degrades progressively before BHEM (red plot in Fig. 4(c)) and remains faulty after retraining (blue in Fig. 4(c)). Degradation is accentuated the latter that BHEM takes place. There is a time window (*T*_*t*_ ≲ 300) within which the dominated SOM recovers successfully in more than 50% of attempts (Fig. 4(d)). But this is not strict—successful recovery in delayed BHEM remains high (∼ 40% and fluctuates above 50% repeatedly, even for long *T*_*t*_). A more parsimonious behavior is observed as a function of the dominated map’s quality right after BHEM (Fig. 4(e-f)). There is a clear degradation threshold 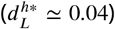 beyond which the likelihood of recovery drops below 50% and vanishes steadily. Hence, in our model, the main contribution towards a critical period does not stem from lost plasticity, but from an ongoing degradation of precursory structures in the dominated side. These structures drift apart because they are idle or adapting to other functions.

## Discussion

The origin of the drastic anatomical transformations after hemispherectomy present a considerable theoretical challenge. BHEM is not only the most remarkable example of brain plasticity: it also offers a unique opportunity to interrogate the origins of the topological mapping between the neural substrate and the emerging representations of input stimuli.

By modeling reorganization of somatosensory cortices, we find two preferred ways in which a healthy topographic representation is reorganized into a smaller, also healthy, map after BHEM:

- **Linear-to-linear** (Fig. 3(a-b)): The intact representations of an optimal linear map are *squeezed* into a smaller region of the neural substrate to make room for orphan stimuli, which are incorporated gradually and in order. This solution appears parsimonious, since the topology of the map is preserved; but it entails a system-wide rewiring of both intact and orphan representations.
- **Linear-to-bilateral** (Fig. 3(c, f-g)): Each orphan, lateralized stimulus is assimilated by the SOM unit that represents its mirror-symmetric counterpart. The topological nature of the map is changed, but the actual rewiring needed is less: intact representations are not displaced and only local adjustments are required.

We prime our networks to help them acquire better representations. This is biologically justified: newborns already have a blueprint of their somatic maps, which mature further as they grow. This blueprint was built over the course of evolution. We find that priming adds a harsh topological constraint: the linear-to-linear reorganization pathway becomes ruled out, but the linearto-bilateral path is very robustly enforced. Furthermore, primed maps conflict with strongly bilateral input stimuli, resulting in pathological representations before BHEM. The robustness of the bilateral attractor after hemispherectomy suggests this intervention to amend defective maps— beyond their current use.

This translates into neuroscientific insights: The lateral-to-bilateral remapping seems a robust, ready-made solution, suggesting why the drastic BHEM has relatively minor cognitive and behavioral impact. Recent fMRI scans indicate that hemispherectomized brains tend to reorganize following this path (***Kliemann et al., 2019, 2021***)—not only for somatosensory networks, but for others such as vision, attention, etc. The same study finds evidence that deviations from this preferred remapping result in worst cognitive performance.

Before BHEM, stimuli with some bilateral symmetry elicit responses in both contra- and ipsilateral areas. Control takes place on the contralateral hemisphere, as activation in the ipsilateral one does not exceed a threshold. In our model, this bilateral activity works as a second priming during training, predisposing far away areas to respond. In the absence of bilaterality, the preferred form of reorganization is a gradual shift of intact representations. A conflict between local (following the linear geometry) versus bilateral reorganization has been observed after stroke (***Pascual-Leone et al., 2005***). Immediately after localized lesions, the ipsilateral (unaffected) hemisphere takes over control of an orphaned limb (e.g. a hand). Patients in which the local map around the injury regains control (presumably by locally reintegrating the orphan limb) recover better. If, instead, the ipsilateral cortex keeps control, recovery is less likely. Tellingly, restriction of movement of the healthy limb and transcranial magnetic stimulation of the unaffected hemisphere (both aimed at switching off the extended ipsilateral control) are promising therapies. Note that this case is different from BHEM: only a patch of map needs reorganization, and the conflict arises from two functioning geometries struggling to take over. Our modeling approach (***Carballo-Castro and Seoane, 2022***) and other similar ones should prove very useful in guiding neurorehabilitation (***Maier et al., 2019***) and recovery after less intrusive surgeries (***Fiest et al., 2014***) as well.

The neural basis of language inspired us to explore how two redundant, competing representations fare. Language develops from bilaterally symmetric circuitry at birth (***Galaburda, 1995; Perani et al., 2011; Berwick and Chomsky, 2016***), usually into a fully lateralized set of interrelated centers (***Geschwind, 1972; Catani et al., 2005; Fedorenko and Kanwisher, 2009; Fedorenko et al., 2012; Fedorenko and Thompson-Schill, 2014; Berwick and Chomsky, 2016***). In our model, a failure to break the initial symmetry can prevent both candidates from maturing, thus losing functionality. This is in agreement with clinical evidence that observes a correlation of symmetry in some language areas across hemispheres and language pathology (***Galaburda, 1995; Bishop, 2013***).

After removal of the language-dominant hemisphere, adults tend to lose that capacity for good (***Harrington, 1995***). Young children can regain function by regrowing key structures in the intact, non-dominant side (***Hertz-Pannier et al., 2002; Liégeois et al., 2008; Danelli et al., 2013; Ivanova et al., 2017; Barceló-Coblijn et al., 2017; Piattelli-Palmarini, 2017***). This suggests critical *window periods* (of around six years (***Woods and Teuber, 1978; Woods and Carey, 1979; Marcotte and Morere, 1990; Hertz-Pannier et al., 2002***)) for function recovery. A frequent interpretation is that plasticity wanes as the brain matures, eventually preventing the re-growth of language centers. But counter-examples exist of older children, even adolescents, regaining full or partial language capacity (***Smith, 1966; Vargha-Khadem et al., 1997; Hertz-Pannier et al., 2002; Voets et al., 2006; Liégeois et al., 2008***). It has also been counter-argued that the adult brain retains its plasticity (***Pons et al., 1991; Pascual-Leone et al., 2005***). A complete, convincing alternative explanation of language loss and recovery mechanisms is missing.

Our results support a promising hypothesis. The key would not lie in plasticity, but in a decay of precursory structures present at birth in the dominated side. Language loss would become irreversible once an integrity threshold is crossed. Crucially, in our experiments the decay of dominated structures is irregularly linked to age: some runs see a swift decline, others keep a fine integrity throughout time. This translates in inconsistent time-window periods—as observed in the clinical record. After long pre-hemispherectomy training, successful re-growing of the dominated map becomes less likely, but it is still a non-vanishing possibility. Stochastic fluctuations make it very probable in odd cases. Our hypothesis depends on the existence of an innate (yet not fully developed) language capacity, as proposed by Chomsky and others (***Bickerton, 1990, 2014; Berwick and Chomsky, 2016***), present in both hemispheres.

Our work relies on two key mechanisms: First, stimuli need to be able to reach every cortical center so that, after BHEM, signals can find a path to candidate representations in the opposite side. We propose a *brain-wide broadcast* of certain neural signals, thus aligning our framework with other hypotheses such as the *global workspace theory* of consciousness (***Dehaene, 2014***). Second, candidate representations must compete with each-other (***Kappel et al., 2014***). Our model also concurs with a view that plastic brain reorganization is aided by unmasking of previously existing (yet inhibited) responses and guided by cortical activity rather than subcortical rewiring (***O’Leary et al., 1994; Pascual-Leone et al., 2005***). Despite its minimal set of assumptions, our model provides deep insights about brain reorganization after hemispherectomy, failure thereof, and potential treatment in certain pathological situations. More realistic models based on our premises might allow us to explore these possibilities in clinical setups.

## Methods

Topographic maps in the somatosensory cortex represent body parts, which appear sorted (from the mouth to the toes) along the distal-medial axis of each hemisphere. This parsimonious representation along a line (which we refer to as a *linear* geometry) mimics the simplest, idealized arrangement of body parts along the body itself. A second relevant geometric component is the bilateral symmetry between parts at each side. Our somatosensory cortices have captured these and other geometrical details of our body throughout our evolution, throughout the development of each individual, and by means of plastic responses to distinct stimuli within a lifetime. Fig. 1 summarizes how we model input signals with one or two conflicting geometries, and how SOMs attempt to infer those geometries through unsupervised learning. In the following we lay out the mathematical details.

**Figure 1.**
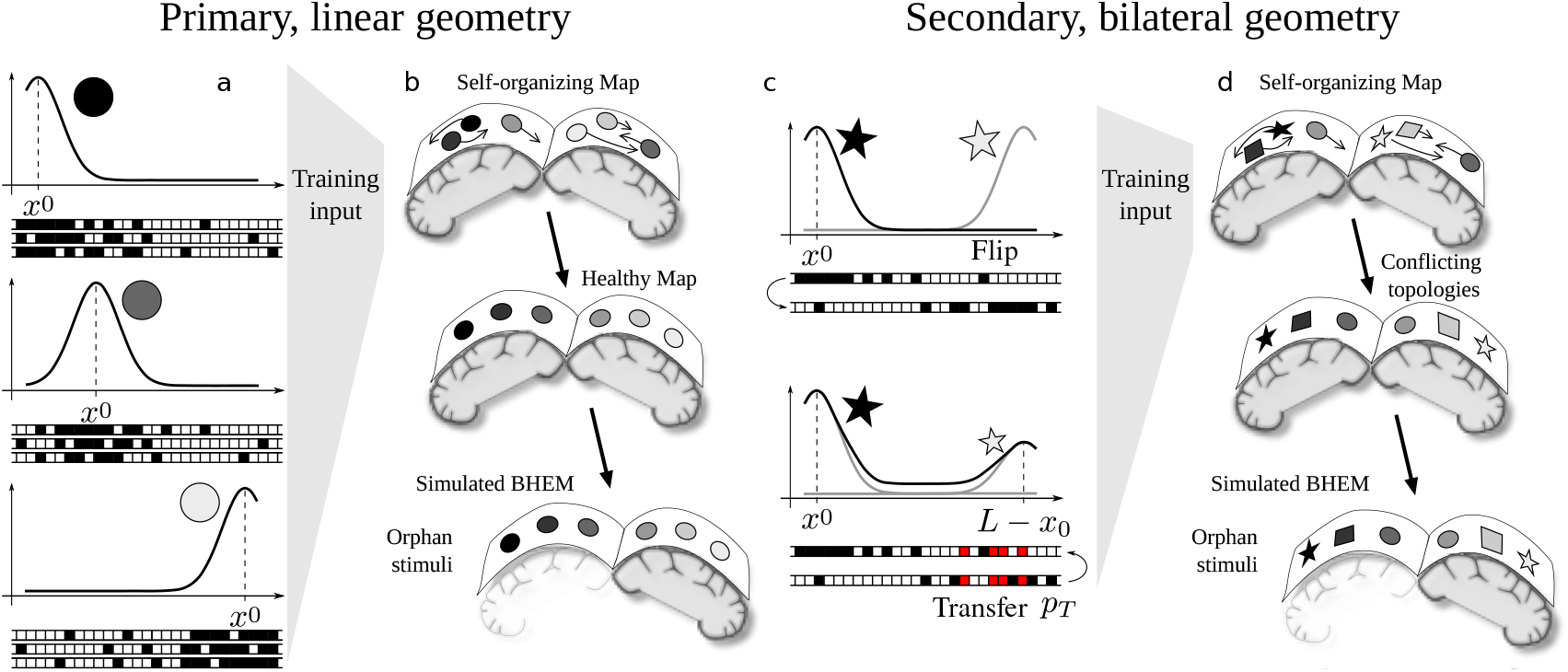
Modeling hemispherectomy with self-organized maps. (a) Each input signal consists of a string of bits (shown under the plots). The curves illustrate the probability that a bit takes value 1 (black boxes in the bit strings). This probability peaks at a different position *x*^0^ ∈ [0, *L*] for each signal, thus giving the range of possible inputs an underlying one-dimensional geometry. A gray gradient (within circles) further illustrates this contiguity. Three possible signals are shown for each *x*^0^ to highlight their stochasticity. (b) Self-organized maps infer the underlying geometry of the data, thus acquiring an internal geometric structure of their own. This geometry might condition reorganization after simulated BHEM (c) We model more complex inputs with the primary linear geometry (gradient-coded) and an additional bilateral symmetry. Therefore we create two bit strings as before, flip one of them, and (with a probability *p*_*T*_) transfer some of the bits of the flipped string (red). This results in signals peaked at a position *x*_0_ and at their mirror-symmetric location (*L* − *x*_0_). The bilateral correspondence is illustrated by geometric shapes, and it conflicts with the simple contiguity of linear inputs. (d) Attempts to learn the conflicting geometries might result in SOMs shifting between alternative representations (Sup. Fig. 3).

**Figure 2.**
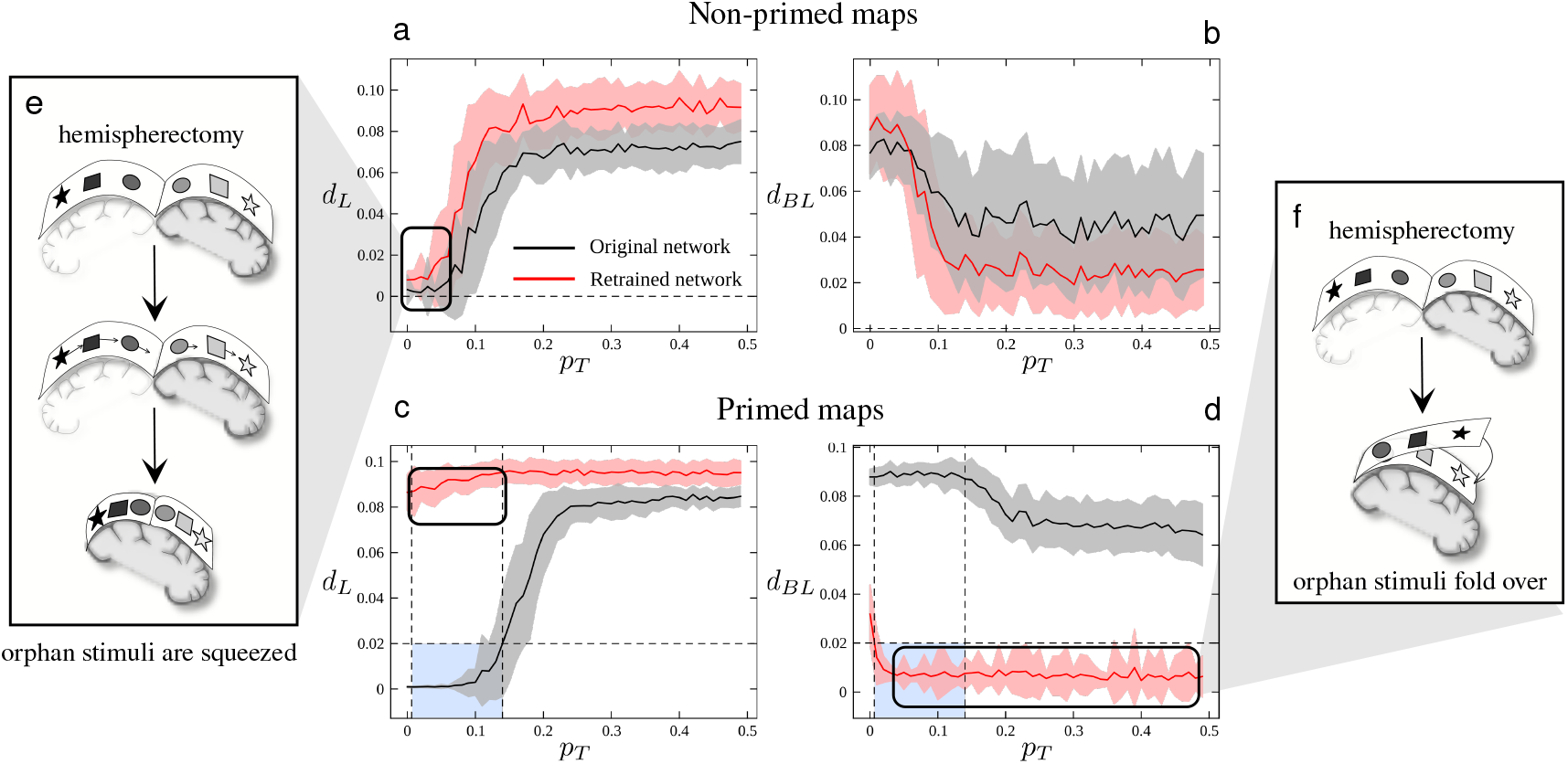
Optimality of emerging maps before and after hemispherectomy. (a-d) SOMs were trained for *T*_*t*_ = 10^4^ steps with signals of fixed bilaterality (*p*_*T*_ ∈ [0, 0.5], horizontal axes). After BHEM, retraining proceeded for *T* ^*rt*^ = 10^3^. Plots display distances to the optimal linear (a, c) and bilateral (b, d) representations right before (black) and after hemispherectomy and retraining (red). Shading indicates standard deviation over 50 repeats. (a-b) Unprimed SOMs with *L* = 100, *σ*_*s*_ = 3, *N* = 200, and *σ*_*t*_ = 30. For low bilaterality, both complete and halved SOMs approach optimal linear representations. Hence, after BHEM, orphan input signals are parsimoniously squeezed into the intact half (e). For higher bilaterality, linear maps fail to form in complete SOMs. Emerging representations approach the optimal bilateral map—even more so in halved SOMs. (c-d) Primed SOMs (*p*_*P*_ = 0.5) with *L* = 100, *σ*_*s*_ = 2, *N* = 100, and *σ*_*t*_ = 10. Optimal linear representations form in complete SOMs and low *p*_*T*_, but not on the halved SOM (black square, very large 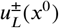). The *squeezing* plasticity pathway is blocked (e). Above a bilaterality threshold, both linear and bilateral representations fail to form in complete SOMs (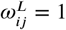 for all *p*_*T*_). But halved, primed SOMs fold very robustly into optimal bilateral representations (f).

**Figure 3.**
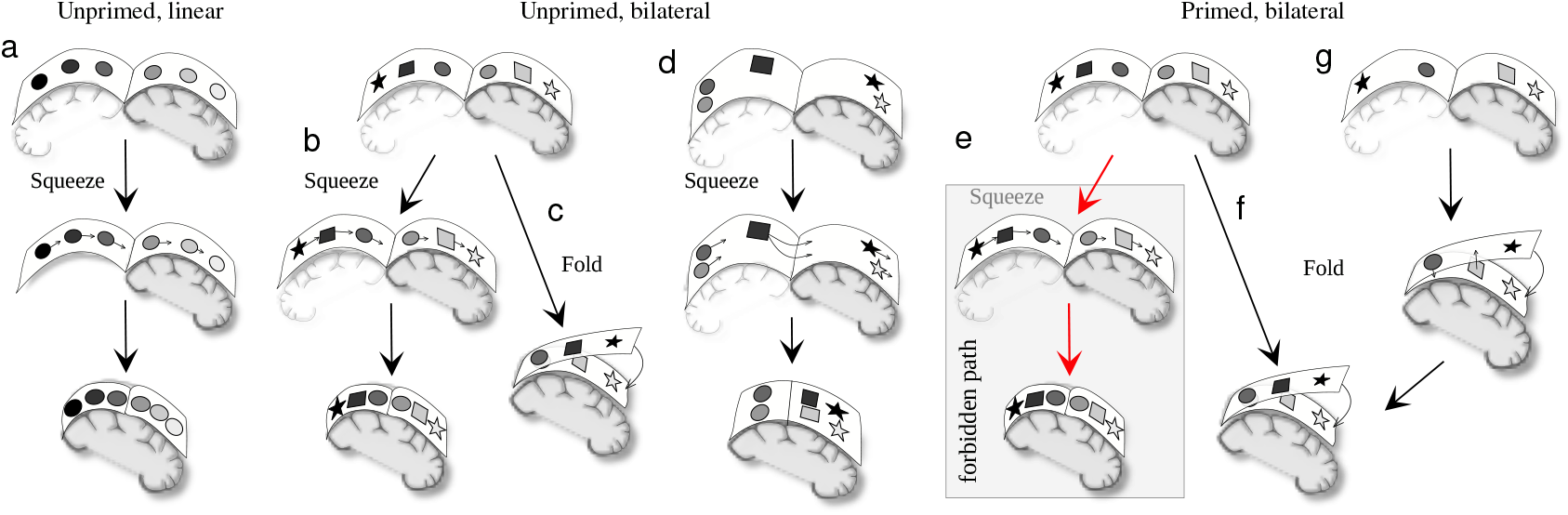
Enabled and forbidden pathways for brain reorganization. (a-b) Reorganization by squeezing orphan stimuli into the intact neural substrate is the preferred form of reorganization for unprimed SOMs and signals with little or no bilateral component. (c) A small fraction (Sup. Fig. 8(a-b)) of primed SOMs trained on moderately bilateral stimuli achieves a linear representation, but after BHEM it is reorganized by folding each orphan stimuli over to its bilaterally symmetric counterpart. (d) Most unprimed, complete SOMs trained on bilateral signals (*p*_*T*_ ≳ 0.1) capture the bilateral (rather than linear) geometry. After BHEM, halved SOMs reorganize into even better bilateral representations. We conjecture that squeezing is also the natural route for this kind of reorganization. (e) The *squeezing* route to reorganization is strictly prohibited for all primed SOMs. (f) In turn, a majority (Sup. Fig. 8(c)) of healthy linear maps fold robustly and swiftly into bilateral representations. (g) Large input bilaterality in primed SOMs results in pathological maps without clear linear or bilateral geometry. This is readily fixed by BHEM, which folds the map into a sturdy bilateral representation.

**Figure 4.**
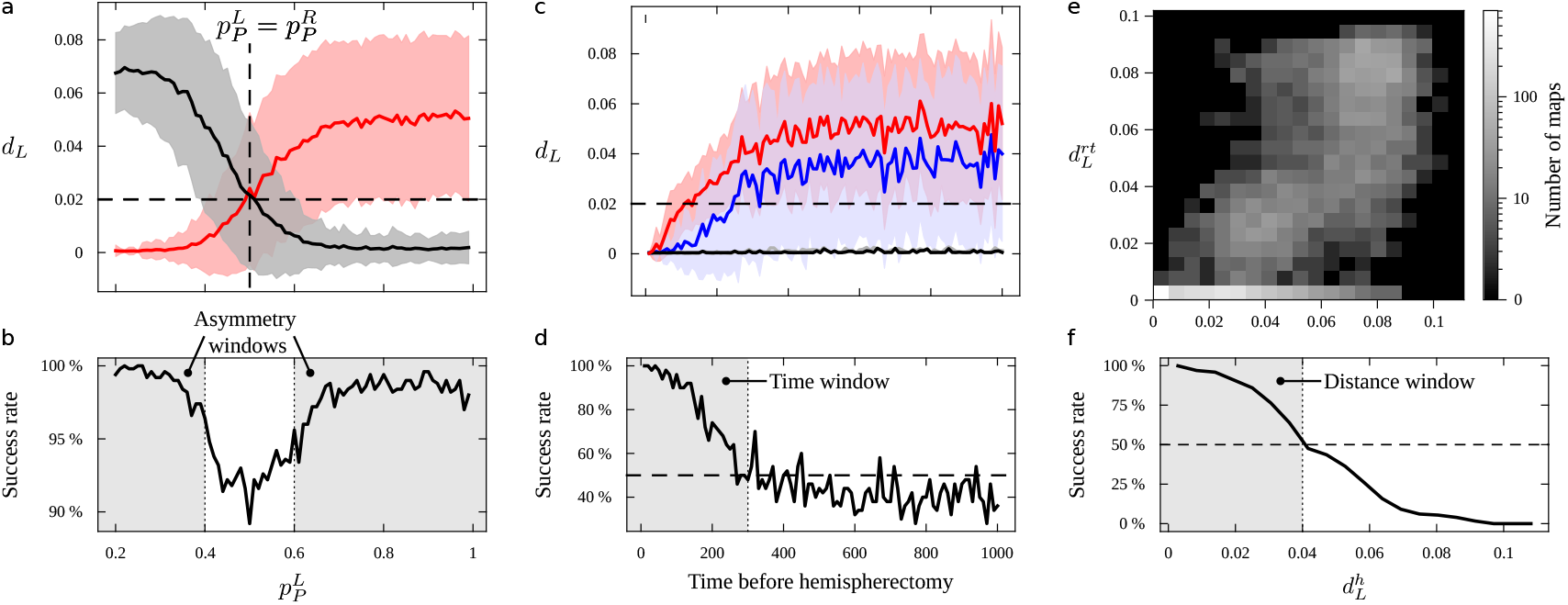
Hemispherectomy in redundant representations. Two half SOMs with *σ*_*t*_ = 10 and *N*_*h*_ = 100 units were concatenated and trained with fully linear (*p*_*T*_ = 0) inputs with *L* = 100 and *σ*_*s*_ = 2. (a) Plots represent distance to the optimal linear map for left (black) and right (red) half SOMs after *T*_*t*_ = 1000 training steps, BHEM, and *T*_*rt*_ = 100 retraining steps. Shading indicates standard deviation over 50 repeats. The right SOM was primed with 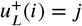 and the left one with 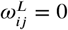 (horizontal axis). (b) Proportion of cases in which a healthy map develops (arbitrarily defined as reaching 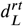). (c-f) We perform BHEM with fixed priming (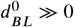 and 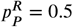, making left dominant) after varying training times *T*_*t*_ ∈ [10, 1000] with *T*_*rt*_ = 100 retraining steps. (c) Plots show distance to optimal linear map for the left SOM before (black) and after (gray) BHEM and retraining (hardly distinguishable), and for the right SOM before (red) and after (blue) BHEM and retraining. Shading (omitted for the left SOM) represents standard deviation over 50 repeats. (d) Proportion of repeats in which the right (dominated) SOM reorganizes into a healthy linear map. (e) Heatmap shows amount of repeats whose right SOM presented a given 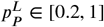 before BHEM (horizontal axis) and reached a given 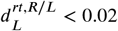 after BHEM and retraining (vertical axis). (f) Proportion of cases that got reorganized into a healthy linear map as a function of 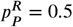 right before BHEM.

**Figure 5.**
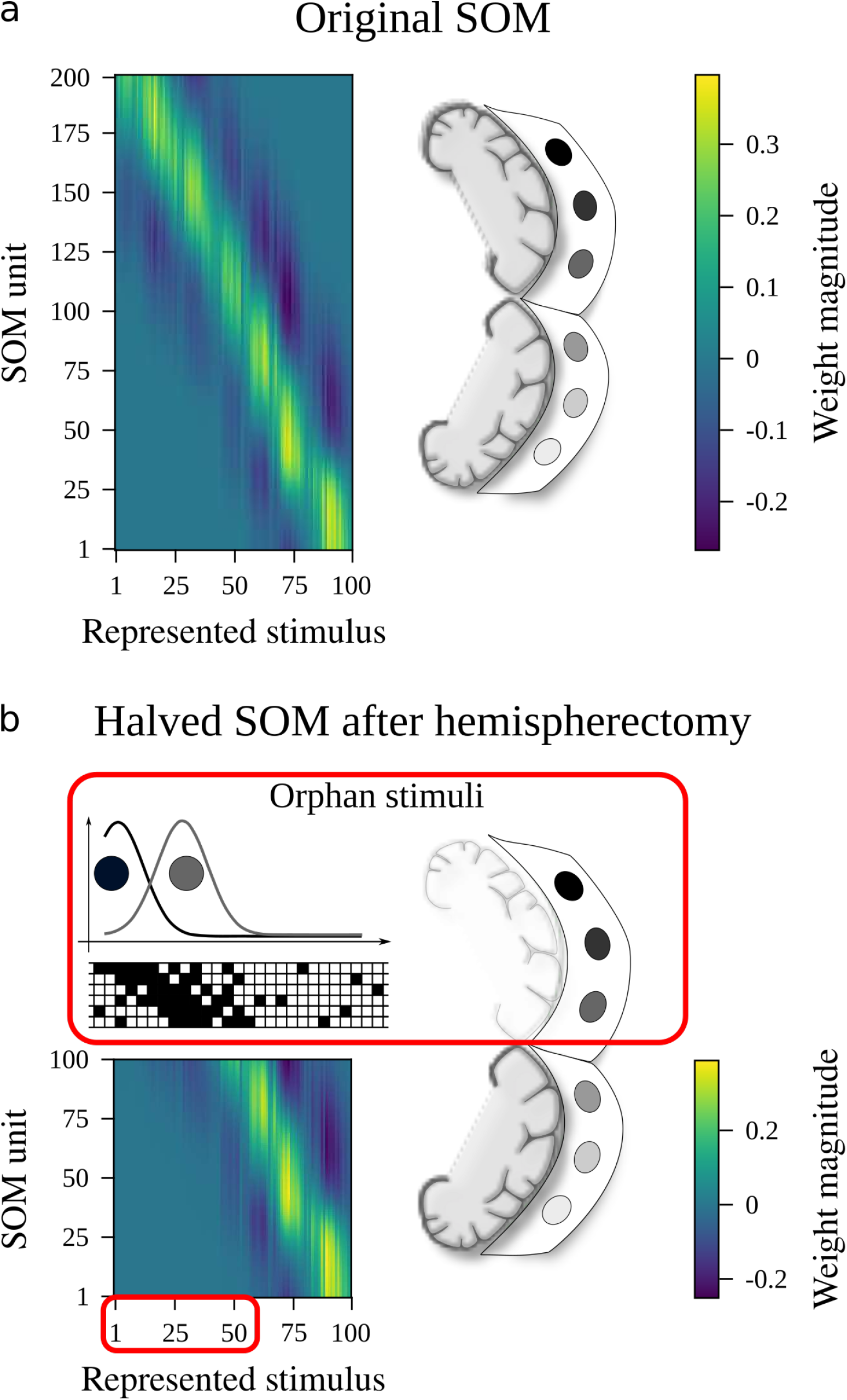
In silico hemispherectomy. (a) A self-organized map infers the geometry of its input signals. An optimal representation of linear inputs results in a roughly linear correspondence between *x*^0^ and the active SOM units, which is also reflected in the SOM’s weights. (b) We simulate hemispherectomy by removing half a SOM. This results in a set of “orphan inputs” whose natural representation is missing.

### Modeling stimuli with linear geometry and bilateral symmetry

Consider a one-dimensional, discretized space of length *L* with boxes centered on *x*_*i*_ ∈ {1, …, *L*}. These boxes can be empty or filled (Fig. 1(a)), thus each contains one bit. The resulting bit string is an input signal, *I*, for our SOM. The value of the box at position *x*_*i*_ is labeled *I*_*i*_ and takes value 1 with probability:

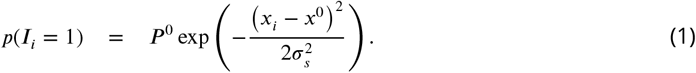

This is a Gaussian normalized by *P* ^0^ = 1 and peaked around *x*^0^ ∈ [1, *L*] with standard deviation *σ*_*s*_. Eq. 1 represents an input elicited at *x*^0^ (think, a touch at a given length along the arm) that has a chance of stimulating areas nearby. The parameter *σ*_*s*_ controls the extent of this spatial correlation. Some stochasticity is present, hence different stimuli centered at *x*^0^ result in distinct binary patterns (each *x*^0^ in Fig. 1(a) is accompanied by three examples). Somatosensory inputs vary over time and originate at changing locations. In our experiments we generate *T* input signals (*I*(*t*), *t* = 1, …, *T*), each peaked at a different (random, uniformly drawn) location *x*^0^(*t*) ∈ [1, *L*].

Real somatosensory inputs are much more complex. Of all the ingredients that enrich our sensory experience, bilateral symmetry plays a key role. Bilaterality is present in the overall structure of the brain (***Harrington, 1995; Swanson, 2000; Seoane, 2020, 2021***) and in somatosensory stimuli—e.g. as both hands engage in a same activity simultaneously. To model a stimulus with bilateral symmetry (Fig. 1(c)), given *x*^0^ we generate two bit strings as above. We flip one of them, thus generating a mirror symmetric stimulus peaking at *L* − *x*_0_. Then, each of the bits that equals 1 in the flipped string is transferred to the unflipped one with probability *p*_*T*_ . This results in a double-peaked probability for *p*(*I*_*i*_ = 1) with the parameter *p*_*T*_ ∈ [0, 1] controlling the level of bilaterality (Fig. 1(c)).

We refer to the signals without bilateral symmetry as having a *linear* geometry. Signals with some *p*_*T*_ > 0 are referred to as *bilateral*. Two relevant model parameters have been introduced so far: *σ*_*s*_, controlling correlations between nearby input positions; and *p*_*T*_, controlling correlations between mirror-symmetric positions.

### Self-Organized Maps models

Self-Organized Maps (***Kohonen, 1982***) are a class of neural networks able to infer, in an unsupervised fashion, the geometrical structure underlaying a given data set. Different SOM versions have been used to model local plasticity in topographic maps after an injury (***Armentrout et al., 1994; Sutton III et al., 1994; Spitzer et al., 1995; Weinrich et al., 1995; Lytton et al., 1999***). Our implementation is the simplest possible, faithful to the original scheme.

Our self-organized network (Sup. Fig. 1(a)) consists of a set of *N* neurons (labeled *j* = 1, …, *N*) that gather external stimuli. Each unit has *L* weights {*ω*_*ij*_, *i* = 1, …, *L*}, labeled as the positions of bit boxes in our input signals. In general, *L* ≠ *N*. At time *t*, the input *I*(*t*) ≡ {*I*_*i*_(*t*), *i* = 1, …, *L*} reaches all units, thus altering their activity, that becomes:

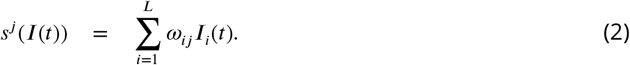

One unit (labeled *k* in Sup. Fig. 1(a)) has a greater activity than any other. We say that it becomes “activated”, while all others are silenced. We say that the active unit *represents* the given stimulus *I*(*t*), and note it:

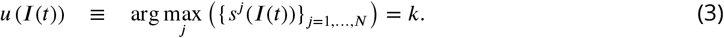

If the network is in training mode, weights are updated according to:

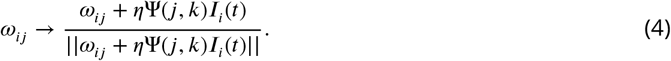

This is, each unit’s weights take a step in the direction of *I*(*t*), thus becoming better at responding to this signal in the future. This step is bounded by a learning rate (here, *η* = 0.1). A further, local modulation is introduced by:

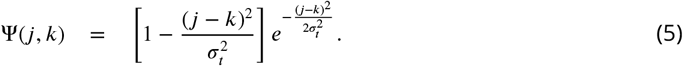

This is the second derivative of a Gaussian centered in the active unit, *k*, and with standard deviation *σ*_*t*_ (Sup. Fig. 1(b)). Ψ(*j, k*) is positive in a neighborhood of *k*. Weights of units in this neighborhood move towards *I*(*t*)—thus also becoming better at responding to this signal. This implements a Hebbian learning step. Outside this neighborhood, Ψ(*j, k*) is negative—thus the corresponding units move away from *I*(*t*) and become worst at responding to it. This implements an anti-Hebbian learning step. Updated weights are normalized to prevent them from blowing up.

These training dynamics introduce a new model parameter, *σ*_*t*_, that encodes spatial correlations between SOM units. These correlations are different from the ones within input signals. In general *σ*_*t*_ ≠ *σ*_*s*_.

### Optimal representations of linear and bilateral stimuli

Eq. 3 defines a SOM’s representation of a specific signal. We are interested in SOM’s responses to classes of signals—specifically, to linear or bilateral inputs as a function of *x*^0^. We denote these representations *u*(*x*^0^) ∈ {1, …, *N*} (with *x*^0^ ∈ [1, …, *L*]), and evaluate them empirically by generating inputs peaked at *x*^0^ and finding the active units as this parameter changes.

A good map will capture the signal’s intrinsic geometry (whether linear or with bilateral symmetry). Hence, the best representation of fully linear stimuli traces a straight line over the space of active SOM units as a function of *x*^0^ (Fig. 5(a)). Two equally good representations exist:

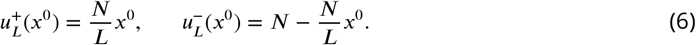

The active SOM unit is a discrete variable, so we can at best approximate 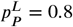. We compute how much does a representation deviate from the closest optimum:

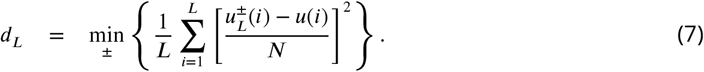

Good representations are i) exhaustive and ii) globally coherent. The first property means that as much space over *x*^0^ ∈ [1, *L*] is covered using as many SOM units as possible. This is achieved if each unit represents a stretch of length *L/N*. But we might arrive to exhaustive representations that are fragmented (Sup. Fig. 2(a-c)), meaning that two nearby positions along *x*^0^ ∈ [1, *L*] are mapped to units which are far appart in the SOM—i.e. there is some *x*^0^ such that ||*u*(*x*^0^)−*u*(*x*^0^ +*ε*) || ≫ 1 for *ε* → 0. In globally-coherent maps, instead, *u*(*x*^0^) changes monotonously as *x*^0^ varies (Fig. 5(a) and Sup. Fig. 2(d-f)).

Optimal representations of stimuli with full bilateral symmetry (*p*_*T*_ = 1) respond equally to inputs peaked at *x*^0^ and *L* − *x*^0^ (i.e., to stimuli peaked at a certain location and to their mirror image). Again two optimal functions solve this problem:

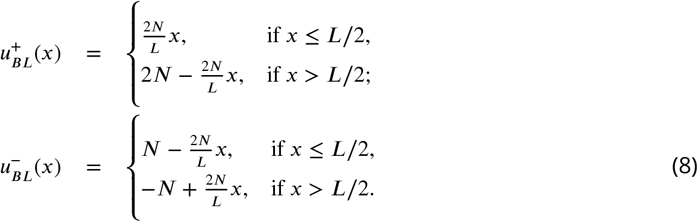

These are piece-wise straight functions shaped into a ‘V’ (Sup. Figs. 3 and 4(d-f)). A SOM’s distance to its closest optimal bilateral representation reads:

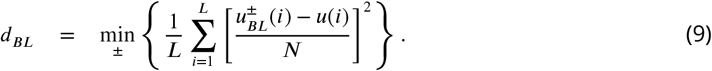

To evaluate a SOM’s distance to the bilateral representation, we activate the network with stimuli *without* bilateral symmetry to ascertain that it is responding to each component separately.

### Priming globally consistent, yet detailed representations

The representations seen so far limit the input and SOM sizes that we can explore. If *L* or *N* become too large, the fixed *σ*_*s*_ and *σ*_*t*_ eventually look small. Then, correlations within inputs or between units are not enough to establish a global coherence in the emerging representations, resulting in fragmented maps. On the other hand, if *σ*_*s*_ and *σ*_*t*_ are kept large, each unit’s response to input stimuli becomes very broad and ambiguous, overlaps with the representations of other units, and the emerging maps do not exploit all resources optimally (Sup. Fig. 5). Ideally, we would like to work with small *σ*_*s*_ and *σ*_*t*_ that would show fine-grained representations, yet still produce globally coherent maps.

To solve this we *prime* our networks with a hint of the correct representation for linear stimuli. So far, SOMs are initialized with random weights *ω*_*ij*_ ∈ *ω*^*R*^, where *ω*^*R*^ ∈ [−1, 1] is a stochastic, uniformly distributed variable. Right after generating them, the *ω*^*R*^ are normalized as in Eq. 4. Now consider a set of target weights *ω*^*L*^ such that 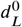 if 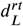 and 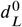 otherwise. This embodies one of the optimal representations in Eq. 6. By initializing our SOMs with:

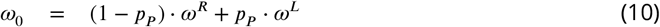

(with *ω*^*R*^ properly normalized before performing this addition), the parameter *p*_*P*_ trades between unprimed and fully primed SOMs. As *p*_*P*_ increases, a situation with fragmented representations (due to small *σ*_*s*_, *σ*_*t*_) transits into a globally coherent map (Sup. Fig. 5). Since we keep *σ*_*s*_ ≪ *L* and *σ*_*t*_ ≪ *N*, a unit’s response does not become coarse.

## Acknowledgments

The authors thank the Complex Systems Lab members for fruitful discussions. Special thanks to Alvah Bassie for his inspiring ideas. This work was supported by the Spanish Ministry of Economy and Competitiveness, grant FIS2016-77447-R MINECO/AEI/FEDER, and an AGAUR FI 2018 grant. Seoane was funded by the Spanish National Reseach Council (CSIC) and the Spanish Ministry for Science and Innovation (MICINN) through a Juan de la Cierva fellowship (grant IJC2018-036694-I) and by the Institute of Interdisciplinary Physics and Complex Systems (IFISC) through the María de Maeztu Program (grant MDM-2017-0711), and by the Jesús Serra Foundation (grant FJSCNB-2022-12-B). This work was done in the Fall of 2019 at the Santa Fe Institute, where the magic happens.

## Supporting Material

**Appendix 0—figure 1.**
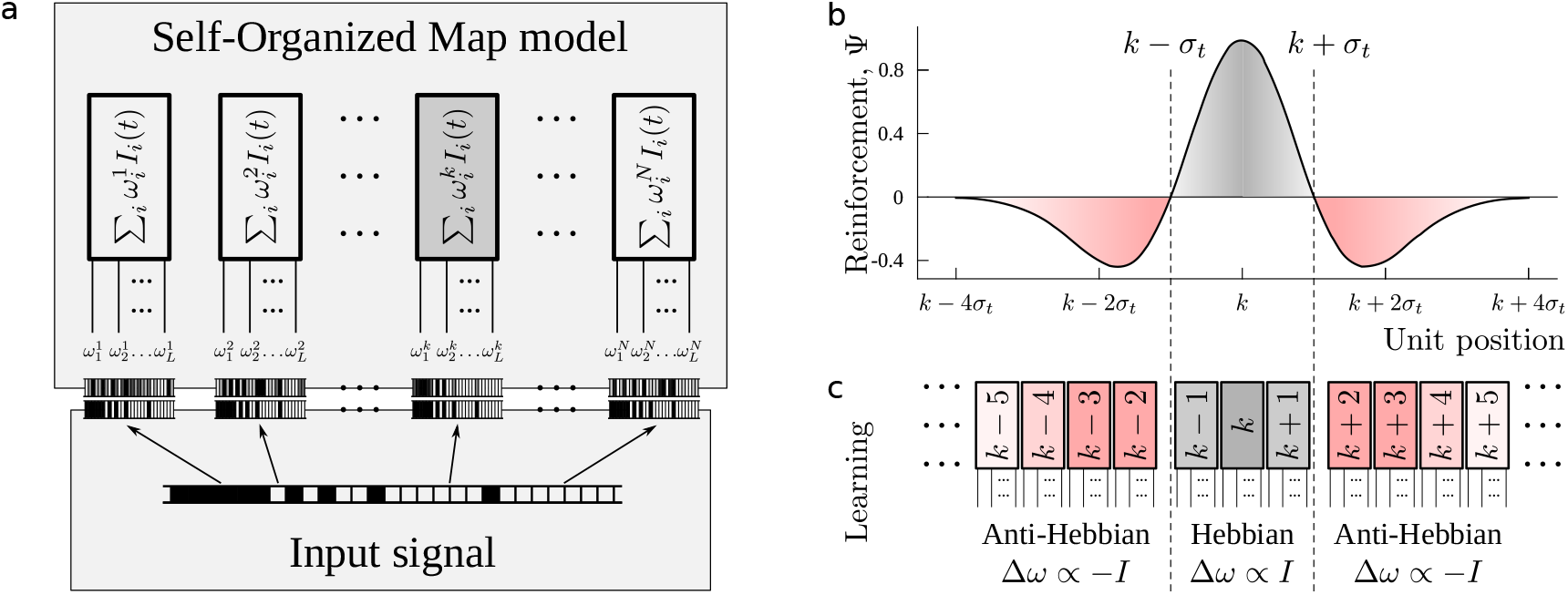
Self-Organized Maps. (a) A one-dimensional binary input is presented to all units of a SOM. Each unit has a set of weights, and one of these sets matches the input signal better than the others (shaded unit, labeled *k*). This unit becomes active and the others, inactive. (b, c) If the SOM is in learning mode, the active unit (*k*) and others in a neighborhood will change their weights in the direction of the current input. Units beyond the neighborhood will updates their weights in the opposite direction. The increase in each unit’s weights is given by a Mexican hat function with width *σ*_*t*_

**Appendix 0—figure 2.**
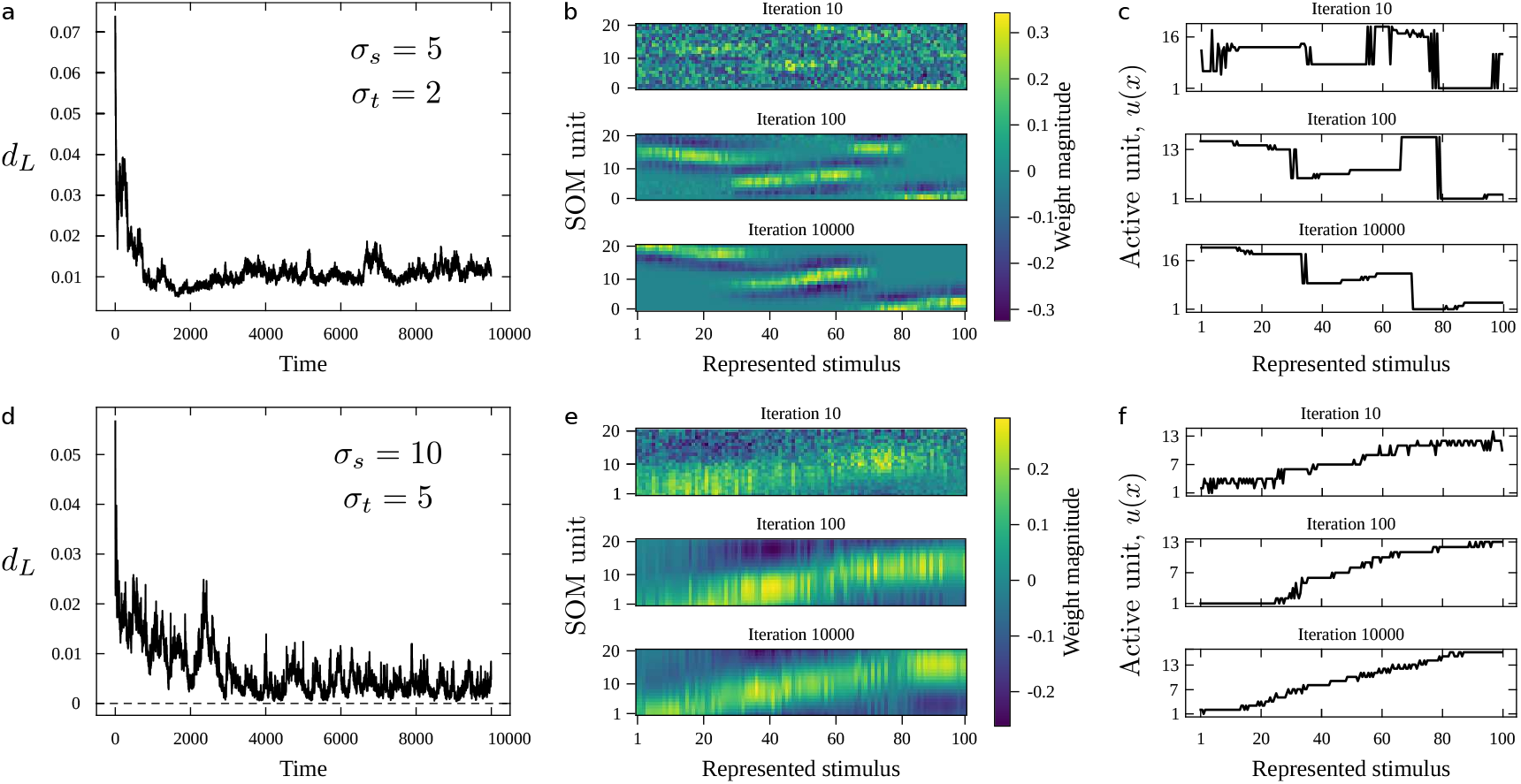
Training Self-Organized Maps for *T*_*t*_ = 10^4^ iterations. (a-c) Convergence to a fragmented representation for *L* = 100, *N* = 20, *σ*_*s*_ = 5, and *σ*_*t*_ = 2. (a) Performance over time, as measured by the distance of the emerging representation to an optimal linear map *d*_*L*_. Average performance stalls at above *d*_*L*_ ≃ 0.01. (b) These heatmaps show the weights for each SOM unit. The horizontal axis represents position in the stimuli space (*x*^0^ ∈ [0, *L*]). The vertical axis represents units in the SOM. The small size of the reinforced neighborhood during training (*σ*_*t*_ = 2) favors that several good representations are nucleated simultaneously along the extent of the SOM. These locally good representations become difficult to flip into a globally coherent one, hence the SOM fails to converge to an optimal solution. (c) Those weights result in a unit that is active when the proper input signal is shown. Fragmentation in the weights results in active units far apart across the SOM for stimuli drawn from very close positions along the stimulus space. (d-f) Convergence to a fine representation for *L* = 100, *N* = 20, *σ*_*s*_ = 10, and *σ*_*t*_ = 6. (d) Performance over time falls below *d*_*L*_ ≃ 0.01, resulting in more optimal representations than the previous case. (e) This is possible due to the coherence brought about by larger correlations both within input signals and between SOM units. (f) Eventually, the SOM representation varies monotonously as stimuli shift smoothly over their range.

**Appendix 0—figure 3.**
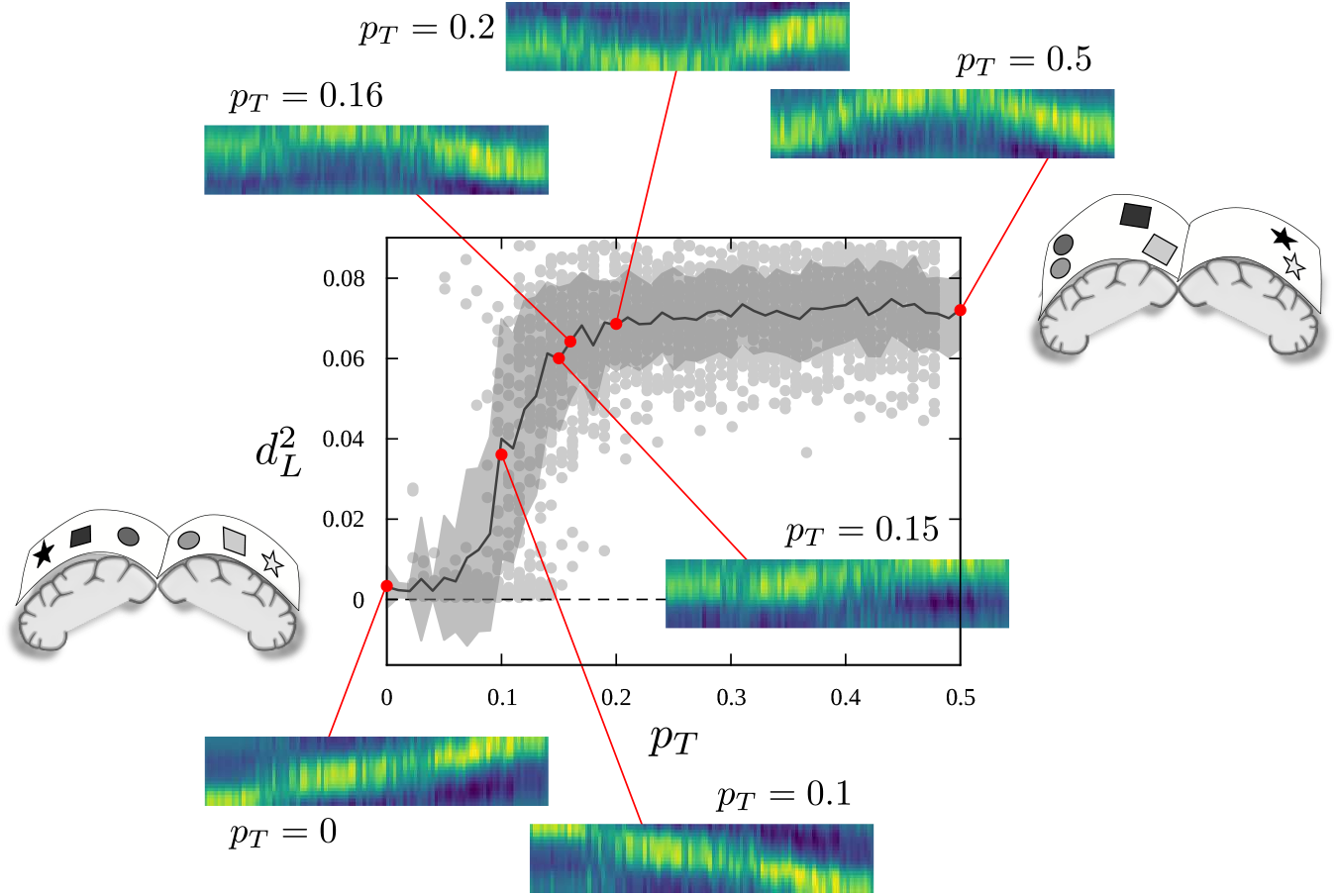
Transitions between representations favoring, respectively, primary and secondary geometries. Maps with *L* = 100, *N* = 20, *σ*_*s*_ = 10, and *σ*_*t*_ = 6 after 10^4^ training steps. SOMs attempt to capture the underlaying geometry of its input signals. When the input can be characterized by a single kind of geometry, SOMs accomodate it more easily than when different geometric symmetries are at play. For low bilaterality in the input, SOMs ignore that component and linear maps emerge. As input bilaterality increases, a conflict arises. Eventually, SOMs ignore the linear geometry and converge to the optimal representation for fully bilateral signals—i.e. each SOM unit responds to signals peaked at some *x*^0^ and to its mirror symmetric signal, peaked at *L* − *x*^0^. Note that this is reflected, for each SOM unit, in weights with sensibility to two mirror symmetric input regions.

**Appendix 0—figure 4.**
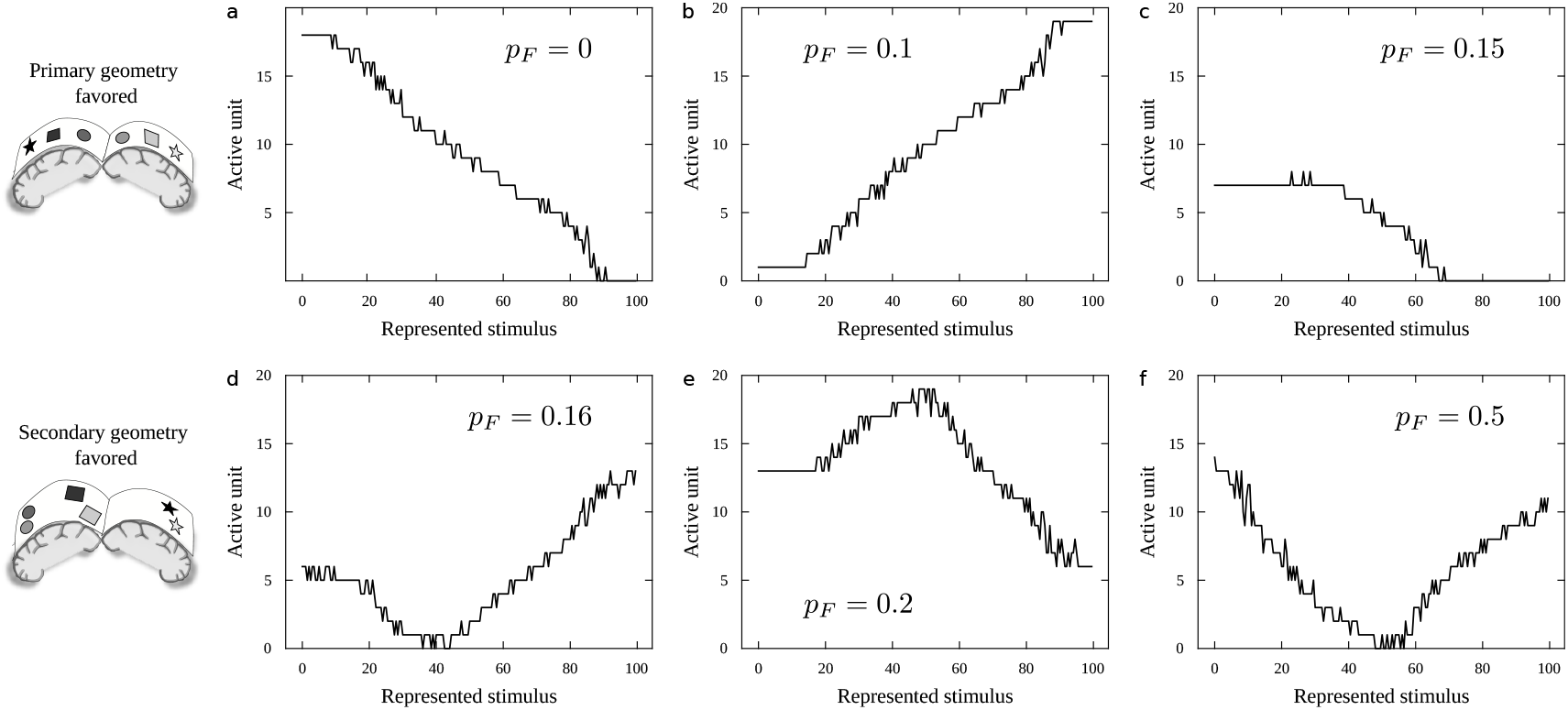
Active units in transitions between primary and secondary geometries. Maps with *L* = 100, *N* = 20, *σ*_*s*_ = 5, and *σ*_*t*_ = 2 after 10^4^ training steps. The weights depicted in Sup. Fig. 3 determine which will be the active unit when a stimulus peaked at a *x*^0^ is presented. These active units follow the geometric hints that the SOM is capturing better. (a-b) The primary, linear geometry is mostly captured, resulting in a linear relationship between *x*^0^ and the index of the active SOM unit. (c-e) At the transition between both geometries, poorer maps emerge. (f) As the bilateral geometry becomes dominant, it is reflected in the active units as well. Again, each unit responds to stimuli peaked at a position *x*^0^ and its mirror symmetric counterpart *L* − *x*^0^.

**Appendix 0—figure 5.**
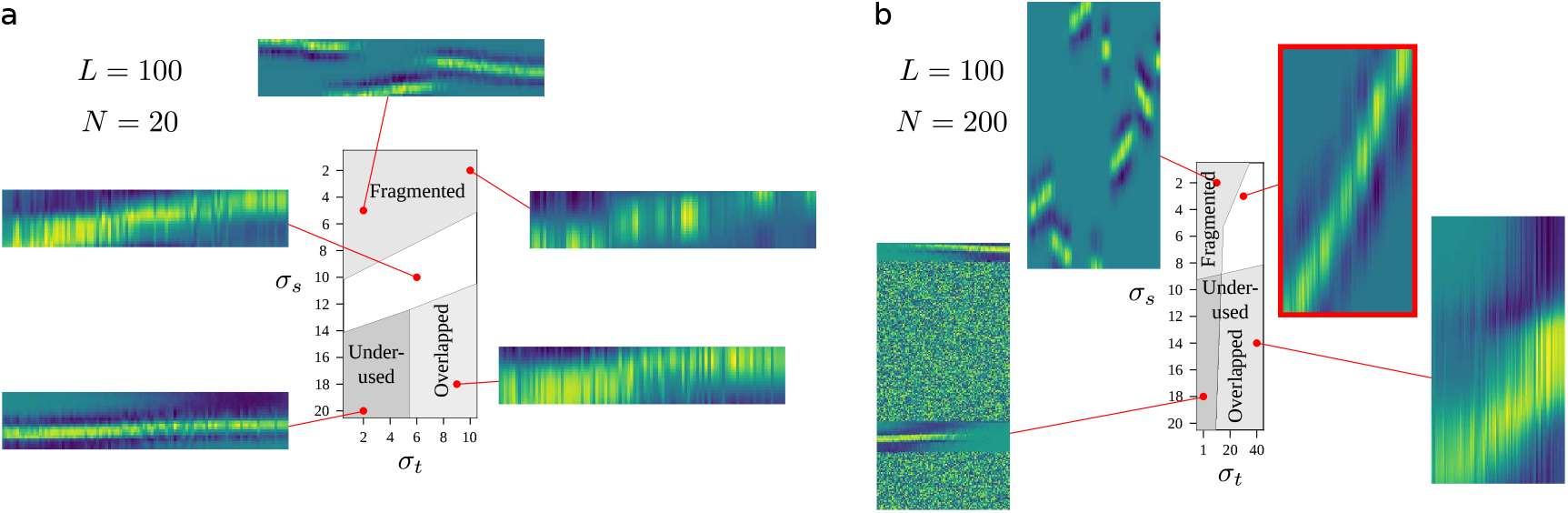
Emerging representations as a function of *σ*_*t*_ and *σ*_*s*_. (a) *L* = 100 and *N* = 20. (b) *L* = 100 and *N* = 200. Relatively small *σ*_*t*_ and *σ*_*s*_ mean that correlations between adjacent input positions or nearby units are short range. This often fails to give the SOM a global coherence and results in fragmented representations. Large *σ*_*t*_ and *σ*_*s*_ can induce global coherence, but at the expenses of generating units with very broad receptive fields that greatly overlap with the receptive fields of other units. If the correlations between units are much smaller than correlations within signals (*σ*_*t*_ ≪ *σ*_*s*_), most units become unused. We generated these maps to inspect visually the quality of the emerging representations. Shaded areas (whose limits are drawn as an illustration) roughly delimit where each kind of faulty representation arises. A balance is possible under the right circumstances—e.g. *σ*_*s*_ = 3, *σ*_*t*_ = 20, framed in red. These model parameters were used for a first round of BHEM simulation.

**Appendix 0—figure 6.**
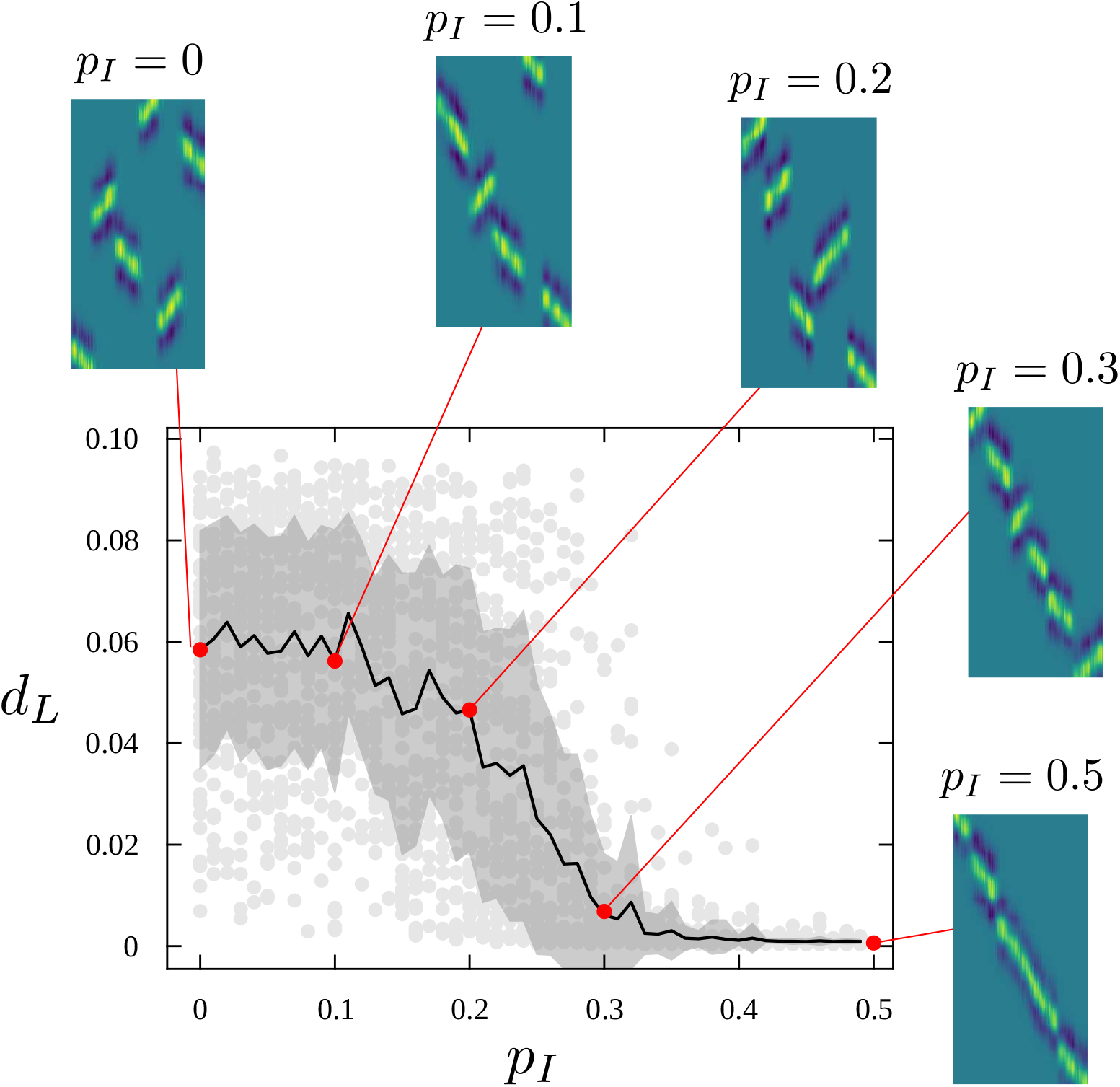
Transition from fragmented to fine-tuned representations. SOMs with relatively low *σ*_*t*_ and *σ*_*s*_ result in fragmented representations because they cannot establish a global coherence (so that multiple, conflicting linear representations start to emerge—but no mechanism exist to align them all in the same direction). Usually, SOMs weights are initialized randomly before training. If, instead, we initialize them with a hint of the global order, a finely-tuned yet global representation emerges more easily. A phase like transition takes us from fragmented representations to globally coherent ones as *p*_*P*_ (the parameter controling the priming of initial weights) increases. Plot shows distance to optimal linear map. Shading represents standard deviation over 50 repeats.

**Appendix 0—figure 7.**
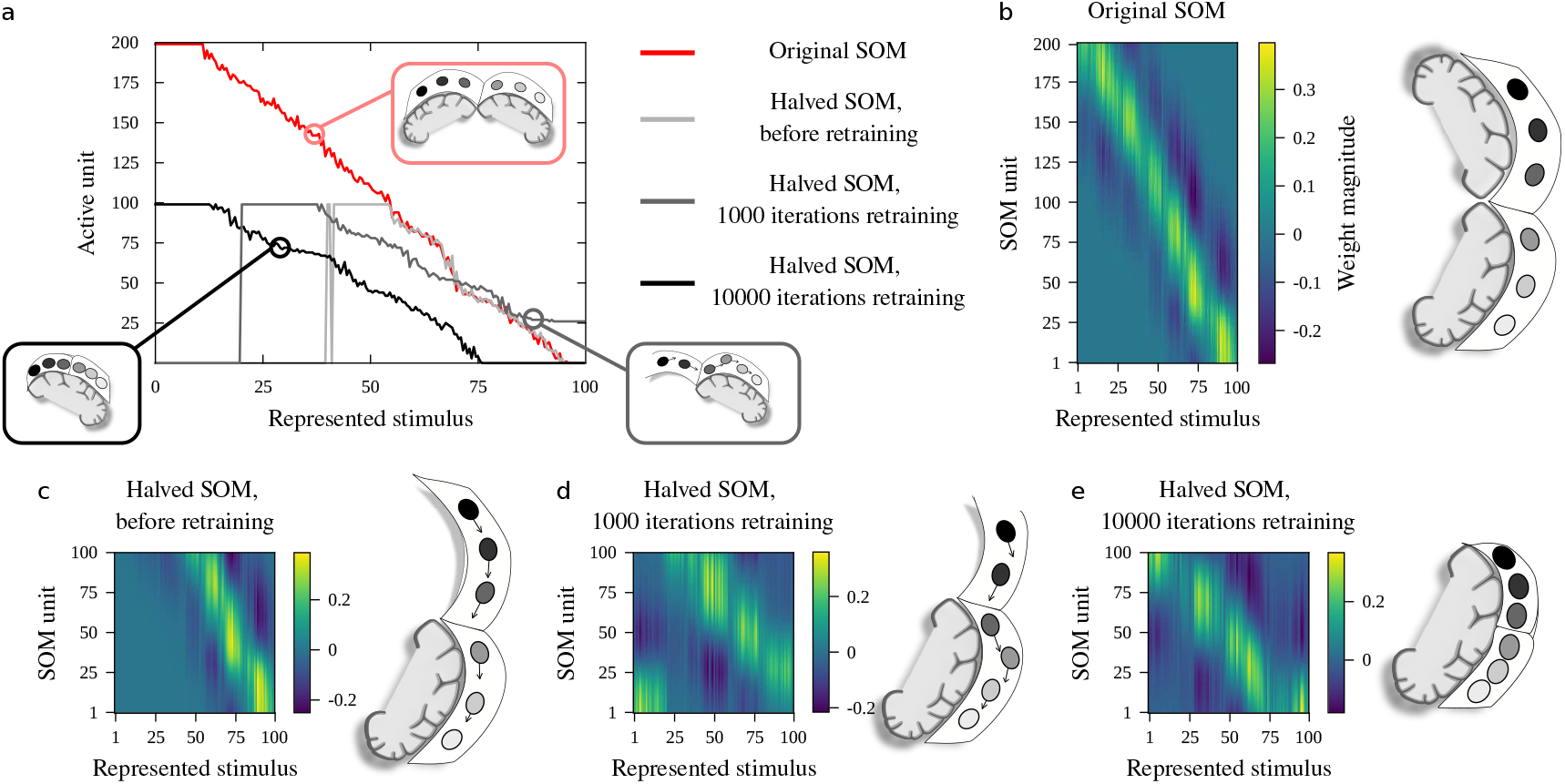
Retraining a SOM to recover a linear representation of the stimuli space. (a) Active units as a function of *x*^0^ in the complete SOM before BHEM (red), and in the halved SOM right after BHEM (light grey), after 10^3^ retraining steps (medium grey), and after 10^4^ retraining steps. (b-e) SOM weights at each of these stages: (b) complete SOM after training, and halved SOM (c) right after BHEM, (d) after 10^3^ retraining steps, and (e) after 10^4^ retraining steps. Each set of weights is accompanied by a cartoon representation of the brain. Note that, in order to keep active units as the variable that depends on the stimuli *x*^0^, the cartoon representation is presented sideways.

**Appendix 0—figure 8.**
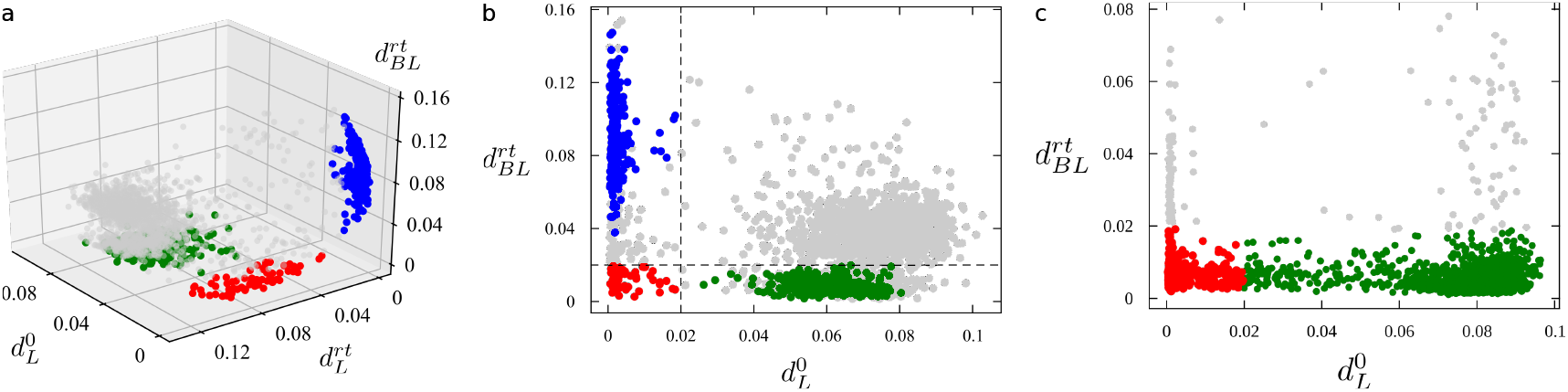
Quantifying what reorganization paths are taken. Each circle represents a BHEM experiment. We depict the distance of the SOM before BHEM to the optimal linear map 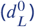 and to the optimal bilateral map after BHEM and retraining 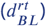. Additionally, in (a) we show the distance to the optimal linear map after BHEM and retraining 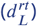. (a-b) Unprimed SOMs with *L* = 100, *σ*_*s*_ = 3, *N* = 200, and *σ*_*t*_ = 30 after 10^4^ training and 10^3^ retraining steps. Gray circles started as linear (low 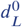) and ended as linear 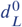 representation, meaning they got reorganized by squeezing. Green started as bilateral representations (low 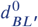, not shown) and ended as bilateral representations as well (low 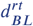). Likely, these maps also reorganized themselves by squeezing—but they ignored the linear geometry from the beginning. Red cases started out as linear representations and ended up as bilateral representation. They got reorganized by folding. (c) Primed SOMs (*p*_*P*_ = 0.5) with *L* = 100, *σ*_*s*_ = 2, *N* = 100, and *σ*_*t*_ = 10. Same training and retraining times. In red, cases that started out as linear and ended up as bilateral representations (i.e. they reorganized themselves by folding). This attractor in halved SOMs is much more stable. Many more cases end up with good bilateral representations disregarding of their starting point (green).

## References

Armentrout SL, Reggia JA, Weinrich M. A neural model of cortical map reorganization following a focal lesion. Artificial Intelligence in Medicine. 1994; 6(5):383–400.

Barceló-Coblijn L, Serna Salazar D, Isaza G, Castillo Ossa LF, Bedia BediaM. Netlang: A software for the linguistic analysis of corpora by means of complex networks. PLoS one. 2017; 12(8):e0181341.

Battro AM. Half a brain is enough: The story of Nico, vol. 5. Cambridge University Press; 2006.

Berwick RC, Chomsky N. Why only us: Language and evolution. MIT press; 2016.

Bickerton D. Language and species. University of Chicago Press; 1990.

Bickerton D. More than nature needs. Harvard University Press; 2014.

Bishop DV. Cerebral asymmetry and language development: cause, correlate, or consequence? Science. 2013; 340(6138):1230531.

Borgstein J, Grootendorst C. Half a brain. The Lancet. 2002; 359(9305):473.

Carballo-Castro A, Seoane SeoaneL. Phase transitions in a simple model of focal stroke imitate recovery and suggest neurorehabilitation strategies. bioRxiv. 2022; p. 2022–12.

Catani M, Jones DK, Ffytche FfytcheD. Perisylvian language networks of the human brain. Annals of Neurology: Official Journal of the American Neurological Association and the Child Neurology Society. 2005; 57(1):8–16.

Danelli L, Cossu G, Berlingeri M, Bottini G, Sberna M, Paulesu E. Is a lone right hemisphere enough? Neurolinguistic architecture in a case with a very early left hemispherectomy. Neurocase. 2013; 19(3):209–231.

Dehaene S. Consciousness and the brain: Deciphering how the brain codes our thoughts. Penguin; 2014.

Devlin A, Cross J, Harkness W, Chong W, Harding B, Vargha-Khadem F, Neville B. Clinical outcomes of hemispherectomy for epilepsy in childhood and adolescence. Brain. 2003; 126(3):556–566.

Fedorenko E, Kanwisher N. Neuroimaging of language: why hasn’t a clearer picture emerged? Language and linguistics compass. 2009; 3(4):839–865.

Fedorenko E, Nieto-Castanon A, Kanwisher N. Lexical and syntactic representations in the brain: an fMRI investigation with multi-voxel pattern analyses. Neuropsychologia. 2012; 50(4):499–513.

Fedorenko E, Thompson-Schill SchillS. Reworking the language network. Trends in cognitive sciences. 2014; 18(3):120–126.

Fiest KM, Sajobi TT, Wiebe S. Epilepsy surgery and meaningful improvements in quality of life: results from a randomized controlled trial. Epilepsia. 2014; 55(6):886–892.

Galaburda AM, Anatomic basis of cerebral dominance. The MIT Press; 1995.

Geschwind N. Language and the brain. Scientific American. 1972; 226(4):76–83.

Harrington A, Unfinished business: models of laterality in the nineteenth century. The MIT Press; 1995.

Hertz-Pannier L, Chiron C, Jambaque I, Renaux-Kieffer V, Moortele PFVd, Delalande O, Fohlen M, Brunelle F, Bihan BihanD. Late plasticity for language in a child’s non-dominant hemisphere: A pre-and post-surgery fMRI study. Brain. 2002; 125(2):361–372.

van den Heuvel MP, Sporns O. A cross-disorder connectome landscape of brain dysconnectivity. Nature reviews neuroscience. 2019; 20(7):435–446.

Ivanova A, Zaidel E, Salamon N, Bookheimer S, Uddin LQ, de Bode S. Intrinsic functional organization of putative language networks in the brain following left cerebral hemispherectomy. Brain Structure and Function. 2017; 222:3795–3805.

Kappel D, Nessler B, Maass W. STDP installs in winner-take-all circuits an online approximation to hidden Markov model learning. PLoS computational biology. 2014; 10(3):e1003511.

Kliemann D, Adolphs R, Paul LK, Tyszka JM, Tranel D. Reorganization of the social brain in individuals with only one intact cerebral hemisphere. Brain Sciences. 2021; 11(8):965.

Kliemann D, Adolphs R, Tyszka JM, Fischl B, Yeo BT, Nair R, Dubois J, Paul PaulL. Intrinsic functional connectivity of the brain in adults with a single cerebral hemisphere. Cell reports. 2019; 29(8):2398–2407.

Kohonen T. Self-organized formation of topologically correct feature maps. Biological cybernetics. 1982; 43(1):59–69.

Liégeois F, Connelly A, Baldeweg T, Vargha-Khadem F. Speaking with a single cerebral hemisphere: fMRI language organization after hemispherectomy in childhood. Brain and language. 2008; 106(3):195–203.

Liégeois F, Morgan AT, Stewart LH, Cross JH, Vogel AP, Vargha-Khadem F. Speech and oral motor profile after childhood hemispherectomy. Brain and language. 2010; 114(2):126–134.

Lytton WW, Stark JM, Yamasaki DS, Sober SoberS. Computer Models of Stroke Recovery: Implications for Neurore-habilitation. The Neuroscientist. 1999; 5(2):100–111.

Maier M, Ballester BR, Verschure VerschureP. Principles of neurorehabilitation after stroke based on motor learning and brain plasticity mechanisms. Frontiers in systems neuroscience. 2019; 13:74.

Marcotte AC, Morere MorereD. Speech lateralization in deaf populations: evidence for a developmental critical period. Brain and Language. 1990; 39(1):134–152.

Merzenich MM, Kaas J, Wall J, Nelson R, Sur M, Felleman D. Topographic reorganization of somatosensory cortical areas 3b and 1 in adult monkeys following restricted deafferentation. Neuroscience. 1983; 8(1):33–55.

Merzenich MM, Nelson RJ, Stryker MP, Cynader MS, Schoppmann A, Zook ZookJ. Somatosensory cortical map changes following digit amputation in adult monkeys. Journal of comparative Neurology. 1984; 224(4):591–605.

Merzenich M, Kaas KaasJ. Reorganization of mammalian somatosensory cortex following peripheral nerve injury. Trends in Neurosciences. 1982; 5:434–436.

Merzenich M, rECANZONE E, Jenkins W, ALLArD T. NUDO, rJ (1988). Cortical representational plasticity. P rakic and W Singer (Eds), Neurobiology of neocortex. 86; p. 41–67.

O’Leary DD, Ruff NL, Dyck DyckR. Development, critical period plasticity, and adult reorganizations of mammalian somatosensory systems. Current opinion in neurobiology. 1994; 4(4):535–544.

Pascual-Leone A, Amedi A, Fregni F, Merabet MerabetL. The plastic human brain cortex. Annu Rev Neurosci. 2005; 28:377–401.

Paul RL, Goodman H, Merzenich M. Alterations in mechanoreceptor input to Brodmann’s areas 1 and 3 of the postcentral hand area of Macaca mulatta after nerve section and regeneration. Brain research. 1972; 39(1):1–19.

Perani D, Saccuman MC, Scifo P, Anwander A, Spada D, Baldoli C, Poloniato A, Lohmann G, Friederici FriedericiA. Neural language networks at birth. Proceedings of the National Academy of Sciences. 2011; 108(38):16056–16061.

Piattelli-Palmarini M. Normal language in abnormal brains. Neuroscience & Biobehavioral Reviews. 2017; 81:188–193.

Pons TP, Garraghty PE, Ommaya AK, Kaas JH, Taub E, Mishkin M. Massive cortical reorganization after sensory deafferentation in adult macaques. Science. 1991; 252(5014):1857–1860.

Pulsifer MB, Brandt J, Salorio CF, Vining EP, Carson BS, Freeman FreemanJ. The cognitive outcome of hemispherectomy in 71 children. Epilepsia. 2004; 45(3):243–254.

Rivera-Rivera PA, Rios-Lago M, Sanchez-Casarrubios S, Salazar O, Yus M, González-Hidalgo M, Sanz A, Avecillas-Chasin J, Alvarez-Linera J, Pascual-Leone A, et al. Cortical plasticity catalyzed by prehabilitation enables extensive resection of brain tumors in eloquent areas. Journal of Neurosurgery. 2017; 126(4):1323–1333.

Seoane LF. Fate of Duplicated Neural Structures. Entropy. 2020; 22(9):928.

Seoane LF. Evolutionary constraints towards lateralization of complex brain functions. arXiv preprint arXiv:211200221. 2021; .

Smith A. Speech and other functions after left (dominant) hemispherectomy. Journal of Neurology, Neuro-surgery, and Psychiatry. 1966; 29(5):467.

Spitzer M, Böhler P, Weisbrod M, Kischka U. A neural network model of phantom limbs. Biological cybernetics. 1995; 72:197–206.

Sutton III GG, Reggia JA, Armentrout SL, D’Autrechy AutrechyC. Cortical map reorganization as a competitive process. Neural Computation. 1994; 6(1):1–13.

Swanson LW. What is the brain? Trends in neurosciences. 2000; 23(11):519–527.

Vargha-Khadem F, Carr LJ, Isaacs E, Brett E, Adams C, Mishkin M. Onset of speech after left hemispherectomy in a nine-year-old boy. Brain: a journal of neurology. 1997; 120(1):159–182.

Voets N, Adcock J, Flitney D, Behrens T, Hart Y, Stacey R, Carpenter K, Matthews P. Distinct right frontal lobe activation in language processing following left hemisphere injury. Brain. 2006; 129(3):754–766.

Weinrich M, Armentrout S, Reggia J. A neural model of recovery from lesions in the somatosensory system. Journal of Neurologic Rehabilitation. 1995; 9(1):25–32.

Woods BT, Carey S. Language deficits after apparent clinical recovery from childhood aphasia. Annals of Neurology: Official Journal of the American Neurological Association and the Child Neurology Society. 1979; 6(5):405–409.

Woods BT, Teuber TeuberH. Changing patterns of childhood aphasia. Annals of Neurology: Official Journal of the American Neurological Association and the Child Neurology Society. 1978; 3(3):273–280.

